# Feed-forward inhibition conveys time-varying stimulus information in a collision detection circuit

**DOI:** 10.1101/217430

**Authors:** Hongxia Wang, Richard B. Dewell, Ying Zhu, Fabrizio Gabbiani

## Abstract

Feed-forward inhibition is ubiquitous as a motif in the organization of neuronal circuits. During sensory information processing, it is traditionally thought to sharpen the responses and temporal tuning of feed-forward excitation onto principal neurons. As it often exhibits complex time-varying activation properties, feed-forward inhibition could also convey information used by single neurons to implement dendritic computations on sensory stimulus variables. We investigated this possibility in a collision detecting neuron of the locust optic lobe that receives both feed-forward excitation and inhibition. We identified a small population of neurons mediating feed-forward inhibition, with wide visual receptive fields and whose responses depend both on the size and speed of moving stimuli. By studying responses to simulated objects approaching on a collision course, we determined that they jointly encode the angular size of expansion of the stimulus. Feed-forward excitation on the other hand encodes a function of the angular velocity of expansion and the targeted collision detecting neuron combines these two variables non-linearly in its firing output. Thus, feed-forward inhibition actively contributes to the detailed firing rate time course of this collision detecting neuron, a feature critical to the appropriate execution of escape behaviors. These results suggest that feed-forward inhibition could similarly convey time-varying stimulus information in other neuronal circuits.

## Introduction

Within neural networks, inhibition operates in close concert with excitation to shape the firing properties of individual neurons. In awake mammalian cortex, it helps define the spatial and temporal spread of activity, promoting sparse firing to sensory stimuli [1, 2]. Inhibition is often subdivided in three types, thought to possess distinct computational properties: feed-forward, lateral and feedback [3]. Feed-forward inhibition enhances the temporal fidelity of single neurons, including Purkinje and pyramidal cells [4-7]. In invertebrates, feed-forward inhibition sharpens the responses of mushroom body Kenyon cells to odors [8], is required for appetitive memory expression [9], and plays a role in motion detection [10]. Lateral inhibition often affects visual responses to spatially extended stimuli, such as those of object-detecting neurons [11]. Here, we focus in a well-defined neurobiological context on a topic that has received less attention: whether feed-forward inhibition conveys detailed time-varying information about spatially extended stimuli beyond its temporal sharpening and synchronizing effects and whether that information could be used to implement specific neuronal computations. In contrast, this issue has been extensively studied for feed-forward excitation.

The lobula giant movement detector (LGMD) in the locust optic lobe is an identified neuron [12], selectively responding to objects approaching on a collision course or their two-dimensional simulation on a screen (looming stimuli) [13-15]. The LGMD receives both feed-forward excitatory and inhibitory inputs and conveys its spiking output to a postsynaptic identified pre-motor neuron, the descending contralateral movement detector (DCMD) [16]. Feed-forward excitation impinges on the largest of three dendritic fields the LGMD possesses and is sculpted by several presynaptic mechanisms: lateral inhibition, global (normalizing) inhibition, and lateral excitation [17-19]. Feed-forward inhibition is subdivided in two distinct channels which sense ON (luminance increments) and OFF (luminance decrements) transients and impinges on two additional dendritic fields [18a, 17, 21]. Feed-forward inhibition helps terminate the LGMD’s response to looming stimuli [21] and interacts non-linearly with feed-forward excitation within the LGMD’s dendritic tree [15, 20, 22], a process thought to be critical to the generation of escape behaviors [23]. However, little is currently known about neurons presynaptic to the LGMD mediating feed-forward inhibition. In the locust medulla, the neuropil upstream of the lobula, non-directional motion-sensitive columnar and tangential neurons that may provide feed-forward inhibition to the LGMD have been characterized [24-26].

To isolate neurons contributing to feed-forward inhibition in this collision detection circuit, we carried out in vivo intracellular LGMD recordings and simultaneous extracellular recordings from putative presynaptic inhibitory neurons. We studied the visual receptive fields and characterized the response properties of those neurons identified to be presynaptic to the LGMD. We found that they have wide receptive fields and exert inhibition through GABA_A_ receptors. Interestingly, their firing pattern was tightly coupled to that of the LGMD, gradually increasing, peaking and decaying towards the time of collision. Because the firing rate of the LGMD is described by multiplying the angular speed of an approaching object with a negative exponential of its angular size, this suggested that feed-forward inhibition might code for angular size [20]. Excitation has already been shown to encode a function of angular speed [27]. The electrophysiological recordings we report allowed us to test whether the firing rate of inhibitory neurons presynaptic to the LGMD encodes the angular size, or speed of looming stimuli, or functions of them. Our results show that feed-forward inhibition conveys to the LGMD time-varying information about an approaching object’s size, in parallel to the speed information conveyed by feed-forward excitation.

## Results

### Feed-forward inhibition mediated by DUB afferents and postsynaptic GABA_A_ receptors reduces the LGMD’s excitability

The LGMD neuron possesses three dendritic fields, labeled A, B and C (Figure 1A and S1A). Dendritic field A receives retinotopically organized, motion-sensitive excitatory inputs from an entire visual hemifield. These inputs are provided by an array of columnar neurons originating at the ommatidia of the compound eye lattice [28, 29]. The axons of columnar retinotopic neurons mediating these inputs form two optic chiasms. The first one is located between the lamina and the medulla (Figure S1B), and the second one between the medulla and the lobula. Dendritic field C, on the other hand, receives inputs thought to mediate inhibition to transient light OFF stimuli [18a]. Anatomical and electrophysiological evidence suggests that these inhibitory inputs are mediated by ∼500 axons originating in the medulla that form an uncrossed axonal tract called the dorsal uncrossed bundle (DUB, Figure S1B) [18a, 30, 31].

**Figure 1.**
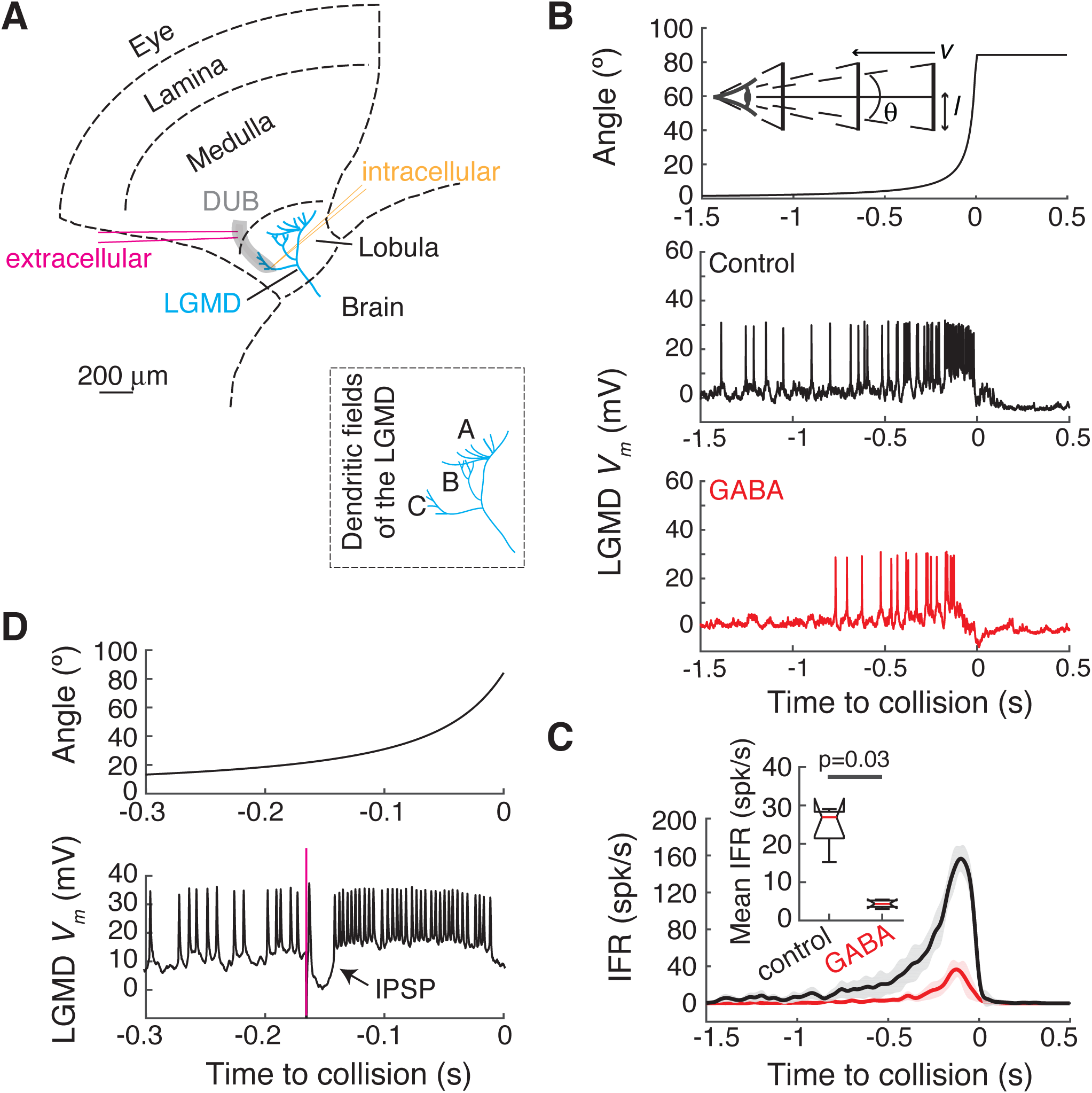
Feed-forward inputs from the dorsal uncrossed bundle to field C of the LGMD are inhibitory. (A) Schematics of the recording configuration in the locust optic lobe. Magenta and dark yellow lines represent the location of the extracellular electrodes and the intracellular one positioned in field C of the LGMD, respectively. Lower right inset shows the three dendritic fields of the LGMD. (B) Intracellular recording from LGMD’s field C (middle, control) in response to a looming stimulus (*l*/|*v*| = 40 ms, top). After puffing GABA on dendritic field C (bottom, GABA), the number of spikes was substantially reduced. 0 is the resting membrane potential. Top diagram: schematic of the stimulus configuration. θ: half-angle subtended at the retina, *l*: stimulus half-size, *v*: approach speed. (C) Mean instantaneous firing rate of the LGMD before and after GABA puffing. Shaded regions denote ±1 s.d. (n = 5 animals). Inset: the mean firing rate averaged over the entire trial was significantly smaller after GABA application (p = 0.03, n = 5, Wilcoxon signed-rank test). For each box plot, the red line shows the median, the upper and lower box limits bracket 75% of the distribution, and the “whiskers” above and below each box show the locations of the minimum and maximum. (D) Electrical stimulation through the extracellular electrodes, (A), during a looming stimulus with *l*/|*v*| = 40 ms (top), triggered a compound IPSP in the LGMD (bottom, arrow). Red line indicates the time of electrical stimulation. See also Figure S1.

To identify DUB neurons presynaptic to the LGMD, we inserted an intracellular electrode in dendritic field C and a pair of extracellular electrodes (stereotrode) ∼200 µm away on the dorsal aspect of the lobula, where the DUB lies (Figure 1A). First, we confirmed that dendritic field C receives inhibitory inputs by puffing GABA close to it and monitoring the membrane potential (*V*_*m*_) of the LGMD in response to looming stimuli. We used stimuli with a ratio *l*/|*v*| = 40 ms, where *l* represents the half-size of the approaching object, and *v* its translation speed towards the locust eye (Figure 1B, top diagram). The responses of the LGMD are depicted in Figures 1B and C. As the simulated object approaches, its angular size (*θ*) increases nearly exponentially and the LGMD firing rate gradually increases, peaks and finally decays towards the time of collision (t = 0), consistent with previous results [20, 21]. GABA application (∼300 µM; Methods) significantly reduced (and sometimes completely abolished) LGMD firing to looming stimuli (Figures 1B and C). Additionally, in the first trial following GABA application, we found the LGMD’s *V*_*m*_ to be tonically hyperpolarized by 2.9 ± 2.4 mV (mean ± s.d., n = 5 animals, average over 1 s before stimulus onset, p = 0.0313, one-sided sign-rank test; mean over five successive trials: 2.1 ± 2.3 mV, p = 0.0625, one-sided sign-rank test). Further, the effects of GABA are consistent with previous findings that the GABA_A_ receptor blocker picrotoxin enhanced the LGMD’s responses to looming stimuli [20-21]. Together, these results indicate that GABA_A_ receptors are involved in mediating feed-forward inhibition to the LGMD.

To ascertain that our extracellular electrodes were positioned in the DUB we applied bipolar electrical stimulation during the presentation of looming stimuli. Electrical stimulation elicited a tightly coupled compound inhibitory postsynaptic potential (IPSP) in the LGMD that transiently inhibited its spiking response (Figure 1D). The membrane hyperpolarization evoked by electrical stimulation amounted to −13.6 ± 3.1 mV and could be detected at an earliest latency of 3.4 ± 0.6 ms (n = 7; Methods). In contrast, we confirmed that without concomitant visual stimulation, electrical stimulation outside the DUB elicited excitatory responses in the LGMD (Figure S1C-E), and, in the DUB, an hyperpolarization following an initial excitatory postsynaptic potential (EPSP; −4.3 ± 1.5 mV, latency: 4.6 ± 0.9 ms, n = 6; Figures S1F and G).

### Identification of presynaptic inhibitory input neurons to LGMD’s dendritic field C

The experimental setup allowed recording of spontaneous activity from several units with variable spike amplitude in the DUB (Figure 2A, top) and simultaneously the LGMD *V*_*m*_ (Figure 2A, bottom). The *V*_*m*_ in field C exhibited spontaneous EPSPs and IPSPs around rest (−63 mV; Figure 2A, middle inset), resulting in a membrane noise of 0.72 mV (average s.d. of *V*_*m*_, n = 10 animals; field A noise s.d. = 1.05 mV [31a]). Since no relation between the extracellular spikes and *V*_*m*_ was apparent, we resorted to spike sorting and spike triggered averaging of *V*_*m*_ to identify presynaptic neurons to the LGMD.

**Figure 2.**
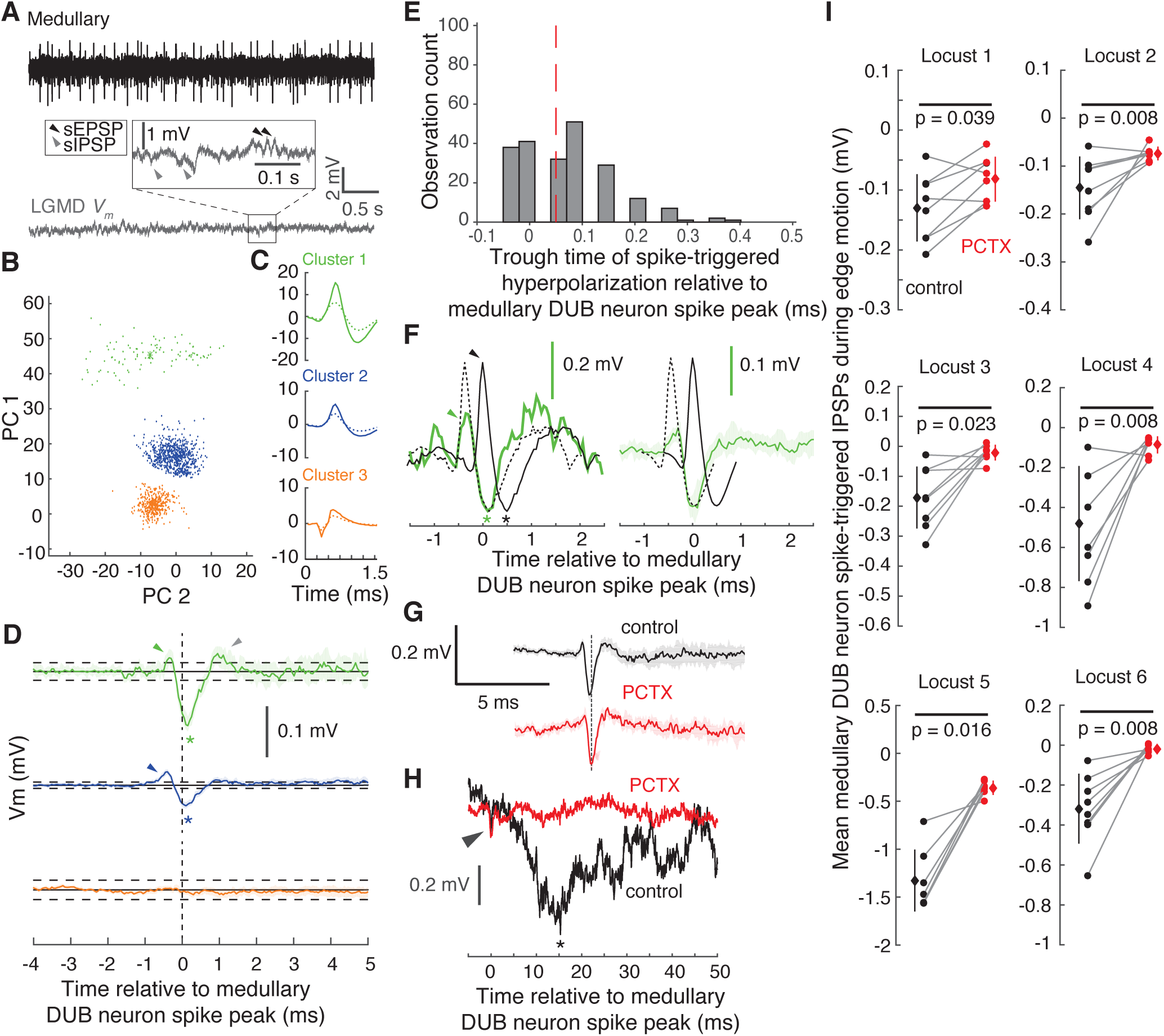
Medullary spike-triggered junction potentials and IPSPs in field C of the LGMD. (A) Example traces of spontaneous multiunit activity of medullary neurons recorded from the DUB (top) and the LGMD’s membrane potential recorded simultaneously (bottom). Inset: close-up view of the LGMD *V*_*m*_. (B) The spikes of medullary neurons were sorted into 3 clusters using PCA and K-means classification. (C) Spike waveforms corresponding to the clusters in (B). The solid and dashed lines correspond to signals recorded at electrode 1 and 2, respectively. (D) Medullary neuronal spikes from clusters 1 and 2 triggered junction potentials consisting of rapid hyperpolarizations of the LGMD *V*_*m*_ in field C, usually preceded by a smaller transient depolarization (green and orange arrowheads). The * symbols indicate the trough of the hyperpolarization. Gray arrowhead: post-hyperpolarization rebound. Dashed lines show the mean ± 4 s.d. of the membrane potential recorded before the medullary spikes. The 3^rd^ cluster spikes (orange, bottom) do not affect the LGMD’s *V*_*m*_. (E) The time to the trough of the spike-triggered hyperpolarization for all the clusters with a significant trough (> 4 s.d. of the mean membrane potential) relative to the time of extracellular medullary spike peak (n = 101 locusts). (F) Left, single spike-triggered hyperpolarization (green) superposed onto the simultaneously recorded extracellular spike triggering it (black). The dashed black line is the same extracellular spike shifted by 0.45 ms. Right, same for the mean spike-triggered hyperpolarization and the mean extracellular spike. Black and green arrowhead: rising phase of extracellular spike and junction potential, respectively. Black and green *: trough of extracellular spike and junction potential, respectively. Note that it this example, the delay from extracellular spike peak (0 ms) to junction potential trough (green star) is 0.15 ms and that this is expected to be an underestimate of the delay between DUB neuron spiking and the LGMD response. (G) Application of picrotoxin (PCTX) onto LGMD’s dendritic field C does not abolish the spike-triggered hyperpolarization. (H) Spike-triggered visually evoked IPSP (*) is abolished by picrotoxin. Arrowhead indicates the spike-triggered hyperpolarization. (I) Reduction of mean spike-triggered visually evoked IPSPs following application of picrotoxin in six animals. The IPSPs were averaged from 5 to 50 ms after medullary spike time peak. Each pair of points represents a different stimulation location on the screen (p-values from Wilcoxon signed-rank test). See also Figure S2.

We performed spike sorting using principal component analysis followed by K-means clustering and identified 3 to 4 clusters per experiment (Figure 2B; see Methods). Spike waveforms consisted of 31 sample points and had different sizes and shapes on the two electrodes (Figure 2C). Next, we performed spike-triggered averages of the LGMD *V*_*m*_ separately for each cluster and determined that some units (typically 2-3) were associated with transient hyperpolarizations of its *V*_*m*_. In Figure 2D, two clusters exhibit such spike-triggered transient hyperpolarizations (Figure 2D, *), while the third cluster did not reveal any associated *V*_*m*_ change above the noise threshold level (Figure 2D, dashed lines). The spike-triggered hyperpolarization events disappeared after randomization of the LGMD *V*_*m*_ relative to the spikes of each cluster, confirming their association with specific clusters (Figure S2A). Additionally, transient hyperpolarizations could be detected in dendritic field C, but not in the excitatory dendritic field. These transient hyperpolarizations, however, were not chemical IPSPs for at least three reasons. First, the duration of the hyperpolarization was short, < 1 ms, unlike GABA_A_ mediated synaptic potentials. Second, they were usually preceded by smaller transient depolarizations (Figure 2D, green and blue arrowheads) and sometimes followed by a rebound (top trace in Figure 2D, grey arrowhead). Third, the latency of their peak (Figure 2D, *) with respect to the extracellular spike peak (dashed blue line in Figure 2D) was short, < 0.3 ms. To confirm this point, we plotted the distribution of trough times across all clusters in all experiments and found a median of 0.05 ms (Figure 2E, dashed red line), within the sampling jitter of our simultaneous recordings (±0.1 ms; Methods). These observations suggested that the spike-triggered transients might be mediated by electrical synapses (gap junctions) and could thus correspond to an attenuated version of the presynaptic neuron’s action potentials.

To investigate this point, we plotted single examples of extracellularly recorded spikes scaled to match the transient *V*_*m*_ hyperpolarization following them (Figure 2F, left) and repeated this procedure after averaging over hundreds of them (Figure 2F, right). We noted that shifting the extracellular spike by ∼0.45 ms yielded a good match between its shape and that of the transient hyperpolarization, both before and after averaging (Figure 2F, dashed lines). Although the extracellularly recorded spike shape may differ from the intracellular waveform, this suggests that the initial rapid phase of the pre-synaptic action potential corresponds to the initial *V*_*m*_ depolarization observed in Figure 2F (black and green arrowheads, respectively) and that this depolarizing junction potential is further attenuated due to its higher frequency content than the following after-hyperpolarization corresponding to the second, slower phase of the extracellularly recorded action potential (hyperpolarizing junction potential; green and black *, respectively), as expected from the low-pass filtering properties of electrical synapses [32]. In invertebrates, gap junctions are formed by innexin molecules distinct from the connexins underlying vertebrate gap junctions [33]. To date no specific blockers are known. Since the vertebrate gap junction blocker carbenoxolone blocks innexins in some cases [34], we probed its effectiveness but found it ineffective against spike-triggered *V*_*m*_ transients (Figure S2B). Hence, to further rule out IPSPs as the source of the transient hyperpolarizations, we applied picrotoxin to dendritic field C and verified that the hyperpolarizing junction potentials remained unaffected (Figure 2G; n = 5 animals).

Next, we reasoned that IPSPs following spontaneous spikes of DUB neurons might be difficult to resolve due to noise in the spontaneous LGMD *V*_*m*_ (Figure 2A). Hence, we stimulated the DUB neurons with small edges moving across the screen (see below for details) and computed spike-triggered averages of the simultaneously recorded LGMD *V*_*m*_. Under these conditions, we could observe an IPSP generated in the LGMD by the DUB neurons (Figure 2H, *). This IPSP disappeared in randomized controls where averaging of the LGMD *V*_*m*_ was uncoupled from the spikes of the studied cluster (Figure S2C). Additionally, the IPSP was nearly abolished by puffing picrotoxin (∼200 µM; Methods) on dendritic field C (Figure 2H, PCTX). Interestingly, the transient hyperpolarization described above could be seen both before and after picrotoxin application further confirming that its origin is unrelated to GABA_A_ receptor activation (Figure 2H, arrowhead). Furthermore, its small size relative to the visually elicited IPSP suggests a negligible role in shaping the LGMD output, in spite of its utility to identify DUB neurons presynaptic to the LGMD.

We repeated this experiment in a total of six animals and computed the mean IPSP elicited by the visual stimulus presented at six locations on the screen before and after picrotoxin application. The size of the visually evoked IPSP varied from animal to animal and from location to location (see below), but picrotoxin reliably reduced it in all animals (Figure 2I). In five of the six experiments, one of the recorded units did not yield a spike-triggered membrane potential hyperpolarization, with a *V*_*m*_ spike-triggered average resembling that of the 3^rd^ unit in Figure 2D (orange trace). In those cases, we also did not find any indication of a spike-triggered IPSP during edge motion. Thus, units that did not exhibit an electrical synapse with the LGMD were also not chemically connected to it.

To summarize, a subset of DUB-recorded neurons provides synaptic input to the LGMD. This input arises both through an electrical and a chemical synapse. The electrical signature of the synaptic connection can be resolved from the spontaneous activity of DUB neurons using spike-triggered averages of the LGMD *V*_*m*_. In contrast, the inhibitory GABA_A_ mediated IPSP cannot be resolved from spontaneous activity but is evident in visually evoked, spike-triggered and averaged activity. This spike-triggered visually-evoked IPSP is largely abolished by picrotoxin.

In total, we recorded from 3-4 DUB neurons per animal across 150 animals. Of those over 60% were electrically connected to the LGMD. Their spontaneous activity was 17.5 ± 12.7 spk/s (mean, s.d.) and was not significantly different for neurons not electrically connected to the LGMD (15.9 ± 6.7 spk/s; p = 0.21, rank-sum test). The peak amplitude of spike-triggered transient hyperpolarizations had a broad range (Figure S2D). As we found that the unit having the largest extracellularly recorded spike was sorted into unit 1, we analyzed unit 1 separately and it consistently generated the largest spike-triggered transient *V*_*m*_ hyperpolarization (Figure S2E, mean amplitude: −0.11 mV, s.d.: −0.04 mV). The median trough latency for unit 1 was also slightly longer than that of other DUB neurons presynaptic to the LGMD (Figure S2F; p = 0.14).

### DUB neurons presynaptic to the LGMD have wide spatial receptive fields

Previous anatomical work suggested that OFF inhibitory inputs to dendritic field C originate from neurons with receptive fields spanning about 8×12° [18a, 30]. To map the receptive fields of the DUB neurons presynaptic to the LGMD, we used OFF edge visual stimuli translating with a constant speed at various locations in the visual field. The edges occupied ¼ of the height (18.8°) or width (22.8°) of the screen and translated at a speed of s28.6 °/s either along the dorso-ventral or along the antero-posterior axis, respectively (Figure 3A, insets). The instantaneous firing rate (IFR) of medullary DUB neurons often exhibited a transient burst immediately after the moving edge had entered the screen (Figure 3A, arrows). After that, the IFR returned close to its spontaneous level (dashed lines in Figure 3A) before tracking the stimulus as it crossed the screen. The strongest sustained responses were obtained for edges moving close to the center of the screen along the dorso-ventral axis, irrespective of motion direction (Figure 3A, top left panel). Responses along the antero-posterior axis were not as sustained (Figure 3A, top right panel) while responses decreased as the stimuli moved closer to the edges of the screen, irrespective of motion direction (Figure 3A, bottom panels). The spatial receptive field obtained by averaging and spatially smoothing the firing rate of 5 DUB neurons with similar receptive fields is shown in Figure 3B. The receptive fields are considerably broader than expected, suggesting the neurons we recorded from are not those characterized in earlier work [18a].

**Figure 3.**
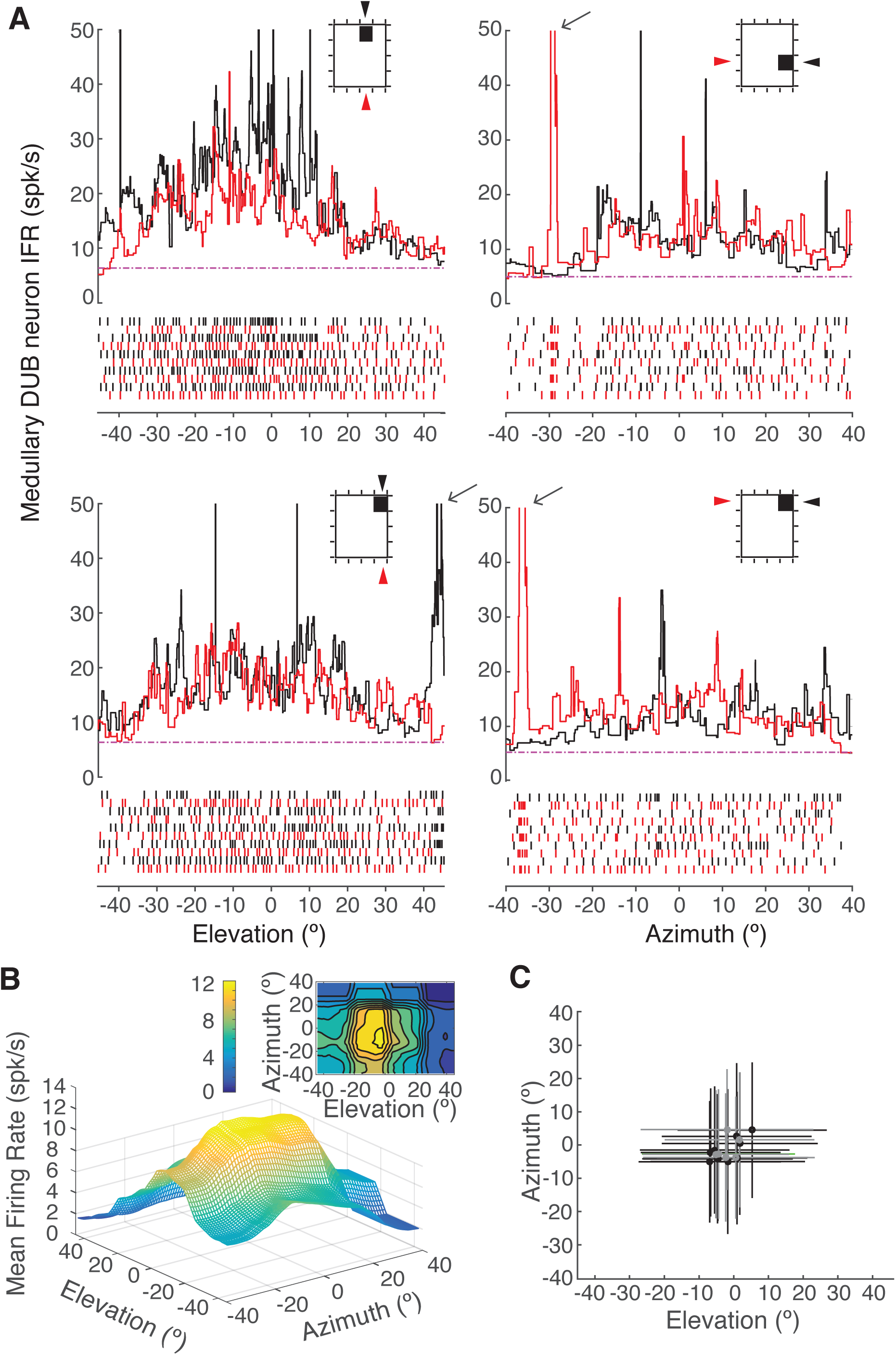
Inhibitory medullary neurons presynaptic to the LGMD have wide receptive fields. (A) Instantaneous firing rate (IFR) responses of medullary neurons to OFF edges translating across the visual field. Left, response of one neuron to 18.8 °-wide edge motion (28.6 °/s) from dorsal to ventral (black) and from ventral to dorsal (red) centered at two azimuth locations. Right, response of a different neuron to 22.8 °-wide edge motion (28.6°/s) from anterior to posterior (black) and from posterior to anterior (red) at two elevations. Insets indicate the direction of motion with arrowheads and the location of the edge on the screen. Point with coordinates (0,0) in elevation and azimuth, respectively, is eye center. Arrows indicate the brief spike burst elicited by the edge appearance on the screen (not always present in plots due to jitter in its occurrence). Bottom rasters are the DUB spikes from 5 trials induced by edge motion. (B) Spatial receptive field obtained by computing the mean firing rate (after subtraction of the spontaneous activity) over five medullary DUB neurons to the stimuli illustrated in (A). Spatial smoothing was achieved by applying a 3×4° mean filter. Upper right inset is the contour plot of the same data. (C) Location of the mean response (black and grey dots) and ±1 standard deviation along azimuth and elevation for 16 inhibitory medullary neurons presynaptic to the LGMD. Black dots indicate cluster with largest spike-triggered hyperpolarization in the LGMD and grey dots other clusters. See also Figure S3.

We also mapped the spatial receptive fields of DUB neurons presynaptic to the LGMD in 5 locusts using edges half as wide. This yielded better spatial resolution at the expense of lower firing rates and thus increased noise. The results resemble those in Figure 3 (Figure S3). Next, we computed the mean location of the spatial receptive field and its standard deviation about the azimuth and elevation axes in 16 DUB neurons presynaptic to the LGMD (recorded in 10 locusts; Methods). Their receptive fields were equally broad as those depicted in Figure 3B (Figure 3C). In contrast, excitatory afferents to dendritic field A of the LGMD have much smaller receptive fields of ∼3° associated with individual ommatidia on the compound eye [29].

### Speed and size tuning of DUB neurons presynaptic to the LGMD

We then examined the dependence of the firing rate of DUB neurons presynaptic to the LGMD on the speed of moving edges. We used 6 speeds varying from 7.1 to 227.2 °/s based on a characterization of LGMD speed tuning [35]. The width of the moving edge was ¼ of the width of the screen, like in the experiments described above. At a speed of 7.1 °/s the OFF-edge stimulus triggered a burst of firing (Figure S4A, arrow) just after entering the screen (left red vertical line). Soon thereafter, the firing rate returned to its spontaneous level and then increased as the stimulus entered the most sensitive part of the neuron’s receptive field. As the edge moved out of the receptive field, the neuron’s firing rate gradually decreased towards its spontaneous level. At intermediate speeds, the burst caused by the stimulus entering the screen was followed by a short period of silence (Figure S4A, 28.4 °/s and 56.8 °/s). At the highest speeds, it merged with the spatial receptive field response (Figure S4A, 272.2 °/s). The peak IFR increased logarithmically with speed, as illustrated across five animals in Figure 4A (dashed lines). In these plots, we pooled data obtained for opposite directions of motion (e.g., dorso-ventral and ventro-dorsal) as no significant difference could be discerned (see below). Speed tuning is weakly dependent on the position of the moving edge along the dorso-ventral axis (Figure 4A left panel) and more strongly along the antero-posterior axis (Figure 4A right panel), with higher responses at higher elevations.

**Figure 4.**
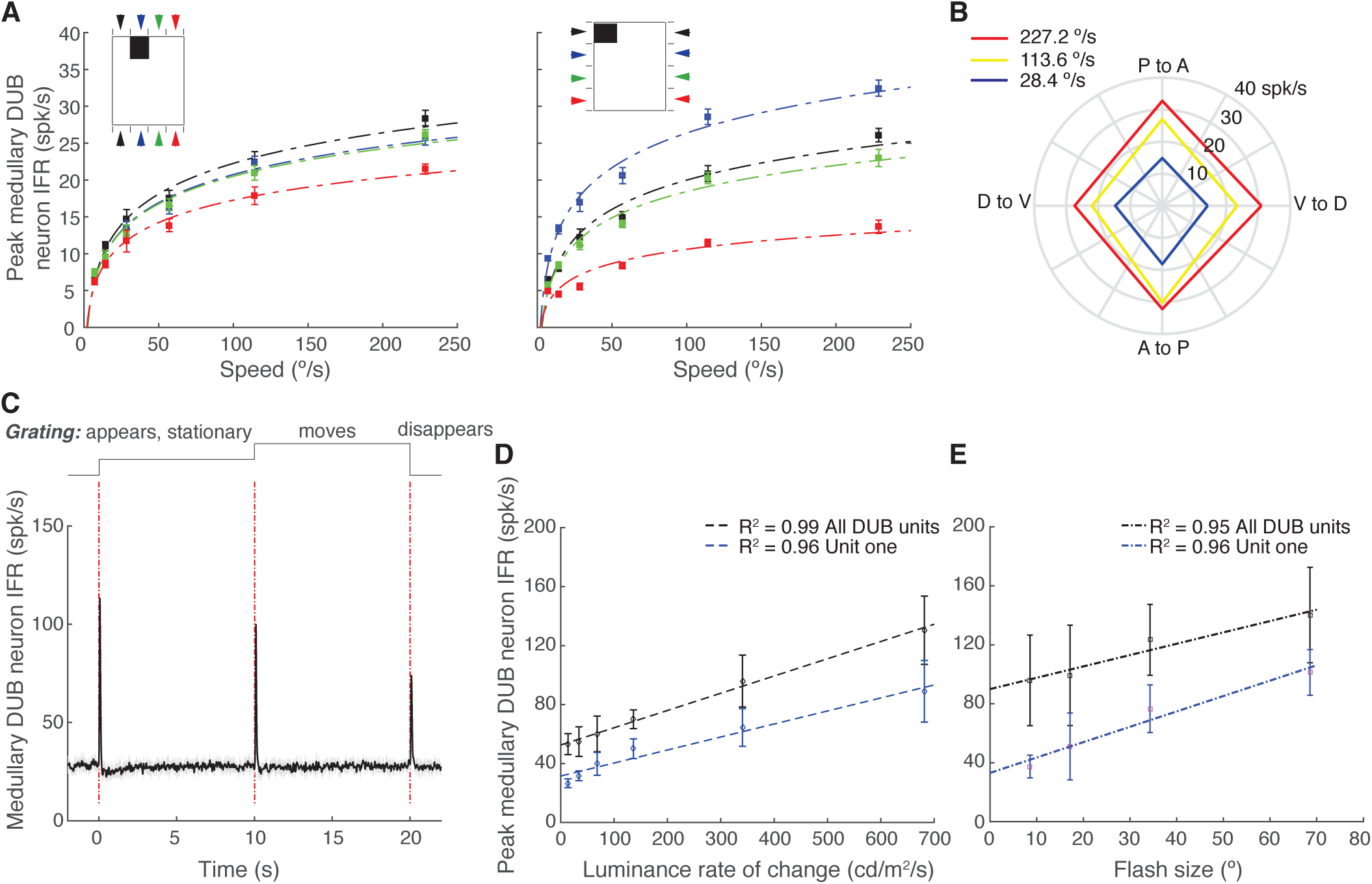
The instantaneous firing rate of medullary DUB neurons presynaptic to the LGMD depends on stimulus speed, size, and rate of luminance change but is independent of movement direction. (A) Peak firing rates of medullary neurons for edges translating from dorsal to ventral and vice versa (left) at four locations on the screen (inset and arrowheads). Right, same as left for anterior to posterior motion and vice-versa (mean ± s.e.m.; n = 5 locusts). Intercepts and slopes from black to red, left: −5.37, −4.65, −4.44, −2.91 spk/s and 6.0, 5.52, 5.42, 4.38 spk/°; right: −6.39, −4.91, −5.22, −2.16 spk/s and 5.73, 6.79, 5.14, 2.77 spk/°. (B) Polar plot of the averaged peak firing rates of medullary neurons across 5 locusts to OFF-edge motion in four different directions. Colors represent different speeds. (C) The IFR of DUB neurons in response to grating movement. Top, timing of stimulus presentation; Bottom black trace is the averaged IFR across 9 locations and 5 locusts. Dashed red lines represent the time at which the grating appears, moves and disappears. Gray edges show 1 s.d. (D) The peak IFRs of DUB unit one (blue) and all DUB units (black) in response to luminance decrement from 68.5 to 0.4 cd/m^2^ over one second, averaged across 5 locusts. Correlation coefficient between luminance change and peak IFR, r = 0.98 ± 0.03 for DUB unit one or 0.96 ± 0.04 (mean ± s.d.). (**E**) The peak IFRs of DUB unit one (blue) and all DUB units (black) in response to OFF flashes with different sizes, averaged across 5 locusts. The goodness of fit, R.^2^ = 0.96 and 0.95 for DUB unit one (blue) and all DUB units. See also Figures S4 – 6.

Additionally, we tested linear models of the relation between peak firing rate and speed. However, we found that in two locations along either the dorso-ventral or antero-posterior direction, log-linear plots fitted better than linear plots (e.g., Figure 4A, left panel, black dashed line, R^2^ = 0.91 ± 0.05 for log-linear and R^2^ = 0.81 ± 0.11 for linear fit, p = 0.0476; blue dashed line, R^2^ = 0.92 ± 0.03 for log-linear and R^2^ = 0.79 ± 0.08 for linear fit, p = 0.0079, one-sided rank sum test). These results resemble those reported for the dependence of the mean firing rate of the LGMD as a function of speed for localized moving stimuli [35]. To confirm the lack of directional selectivity noted in the context of Figure 4A, we used edges moving in four directions close to the center of the receptive field to derive a polar response plot (Figures S4B and 4B). Firing rates in the four directions were nearly identical, and they increased uniformly with speed (results were averaged across 5 neurons).

During these edge motion stimuli, the luminance steadily decreased as the dark edge crossed the screen and the DUB neuron responses increased with edge size (Figure S4C). To test if DUB units are sensitive to motion without luminance change [36], we measured responses to isoluminant gratings at nine different spatial positions in the neurons’ receptive fields. As can be seen in one example cell in Figure S5A, responses were similar at all positions and were thus averaged across spatial locations. The results, depicted in Figure 4C show that there was a transient response to the appearance and disappearance of these gratings as well as the initiation of motion. The latency and strength of response varied slightly for these three distinct events (Figures S5B and C). There was, however, no sustained response to the motion (Figure 4C). Conversely, we tested responses to luminance decrease without any edge motion by changing the whole screen luminance over progressively shorter time intervals. These stimuli elicited transient changes in firing rate that increased in size and decreased in latency as the luminance change intervals became shorter (Figure S6). As demonstrated in Figure 4D, DUB units firing rates increased linearly with the rate of luminance change. In addition, we presented instantaneous luminance changes (flashes), and found that the DUB responses also increased linearly with the size of the flashed region (Figure 4E). This result is consistent with the large receptive fields of DUB units (Figures 3B and C), and the response dependence on edge size (Figure S4C). In summary, no directional selectivity was found among DUB neurons presynaptic to the LGMD. Although their firing rate depended on edge speed, the main determinants of the DUB neurons’ responses appear to be the size and speed of luminance change across their receptive fields.

### The response to looming stimuli of DUB neurons is delayed relative to that of the LGMD

As shown in Figure 1, the firing rate of the LGMD in response to looming stimuli has a characteristic profile. What is the corresponding response pattern of DUB neurons presynaptic to the LGMD? The left panel of Figure 5A illustrates the firing rate of two such neurons (cyan and green traces) recorded simultaneously with the LGMD in response to a looming stimulus with *l*/|*v*| = 20 ms. Both neurons have an activity profile resembling that of the LGMD (red traces in Figure 5A). Interestingly, their firing outlasted that of the LGMD, a characteristic made clearer by pooling their activity (black traces in Figure 5A). Similar results were observed at *l*/|*v*| = 40 ms (Figure 5A, middle panel) and at *l*/|*v*| = 80 ms (Figure 5A, right panel). In all conditions, DUB firing rates always remained elevated for at least 100 ms after the LGMD response stopped before returning to spontaneous firing levels.

**Figure 5.**
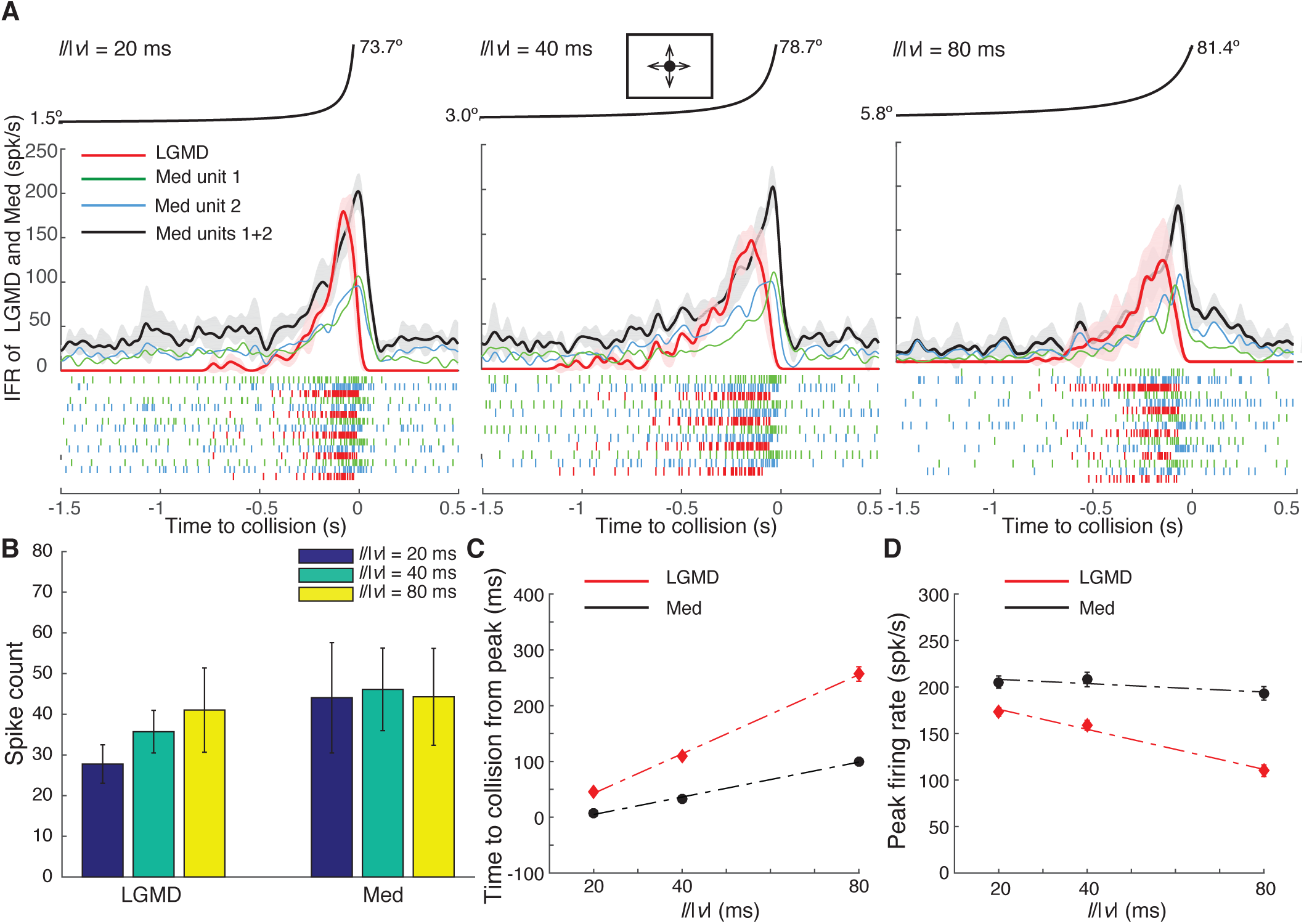
Responses of medullary DUB neurons presynaptic to the LGMD and of the LGMD to looming stimuli. (A) From left to right: example responses to looming stimuli at *l*/|*v*| = 20, 40 and 80 ms (corresponding to three object approach velocities for a fixed object size). Top, the object angular size is plotted as a function of time to collision (t = 0) as the simulated object approaches towards the eye. Middle, instantaneous firing rate of medullary (Med) neurons (green and blue are individual neurons, black is summed across both units) and the LGMD (red) to looming stimuli. Bottom spike rasters indicating the spike times for five trials. Inset shows a schematic of the looming stimuli. Med unit 1 peak times to collision: −8.8 ± 26.9, 40.5 ± 12.4 and 108.0 ± 22.9 ms, respectively (mean ± s.d., n = 7; p < 0.05; one-sided rank sum test). Med unit 2 peak times to collision: −5.5 ± 8.9, 32.6 ± 16.8 and 131.2 ± 112.6 ms, respectively (p = 0.0011 for each paired comparison). Right panel at l/|v| = 80 ms is from a different locust than that used in left and middle panels. (B) Spike count elicited by looming stimuli at the same three *l*/|*v*| values (LGMD: mean ± s.d., n = 7 animals; Med: mean ± s.d., n = 15 neurons in 7 animals). (**C** and **D**) The time at peak firing rate relative to collision and the maximum firing rate for the same three *l*/|*v*| values (mean ± s.d., n = 7 locusts). For medullary DUB neurons, values are reported for summed activity (black lines in A). Dotted lines represent linear fits.

Across our population sample, the spike counts of the LGMD and the DUB neurons were comparable (Figure 5B). The timing of the DUB neurons’ peak firing rate, however, always followed that of the LGMD (Figure 5C). Further, both the DUB and LGMD peak times were tightly correlated with *l/|v|* (Figure 5C) and consequently with each other (ρ = 0.70 ± 0.21, n = 5). The difference in peak firing rate between DUB neurons and the LGMD also increased with *l*/|*v*| (Figure 5D). That DUB neurons respond vigorously to looming stimuli suggests that they contribute to the decay of the LGMD firing rate observed towards the end of a looming stimulus, consistent with changes in LGMD firing rates observed after blocking feed-forward inhibition [21].

We further investigated whether any significant differences could be found between populations of DUB neuron pairs recorded simultaneously. In seven such recordings, we labeled the neuron yielding the largest spike-triggered hyperpolarization in the LGMD as neuron 1, the other one being neuron 2 (c.f. Figure 2B-D, green and blue clusters; Figure 5A). Across experiments, we found that the peak firing times occurred later as the *l*/|*v*| value increased (Figure 5A, legend). However, no significant differences were found for the peak times to collision between the two units at each *l*/|*v*| value (p = 0.12, 0.24, and 0.29 for l/|v| = 20, 40, and 80 ms according to a one-tailed rank-sum test). Additionally, there were no significant differences between the peak firing rates of the two units at each *l*/|*v*| value (p = 0.47, 0.47, and 0.29 for l/|v| = 20, 40, and 80 ms; one-tailed rank-sum test). Thus, the population of DUB neurons presynaptic to the LGMD appears homogeneous with respect to its responses to looming stimuli.

### The instantaneous firing rate of DUB neurons is linearly related to the angular size of looming stimuli

Which features of looming stimuli might be encoded by DUB neurons presynaptic to the LGMD? From earlier work, we know that two variables describe the main characteristics of the LGMD’s firing rate in response to simulated object approach: angular size, *θ(t)*, and angular speed, *θ′(t)* [15, 22]. Pre-synaptic excitatory inputs to the LGMD encode the square of angular speed in their population response [27]. Since, as shown above, DUB neurons modulate their responses both to moving edge speed and size (Figure 4), and since these two variables are correlated during looming, a direct fit of responses to looming stimuli is needed to clarify this issue. We thus plotted the pooled firing rates of DUB neurons as a function of the angular size (Figure 6A; Figure S7A) and speed (Figure S7B) of looming stimuli. For all *l*/|*v*| values, we used data starting 4 s before collision, when the stimulus is small (< 2.3°; Figure 5A), to the end of stimulus expansion. During faster looming stimuli (*l*/|*v*| = 20 ms), the medullary IFR initially increased with angular size and then saturated. However, with higher values of *l*/|*v*|, 40 or 80 ms, this saturation did not occur. For stimuli with *l*/|*v*| = 20 ms, the angles for which the firing rate saturates (>30°) correspond to only the last 50 ms of expansion. This short time, paired with a 20 ms smoothing of the firing rates contributes to the observed saturation. This observation is supported by the improved performance of the angular size model in the time domain (below). The DUB neurons’ firing also increased in response to angular speed at *l*/|*v*| = 20 ms, but rapidly reached a peak and saturated. Saturation was also evident at higher *l*/|*v*| values. These results suggest that DUB neurons encode angular size rather than speed.

**Figure 6.**
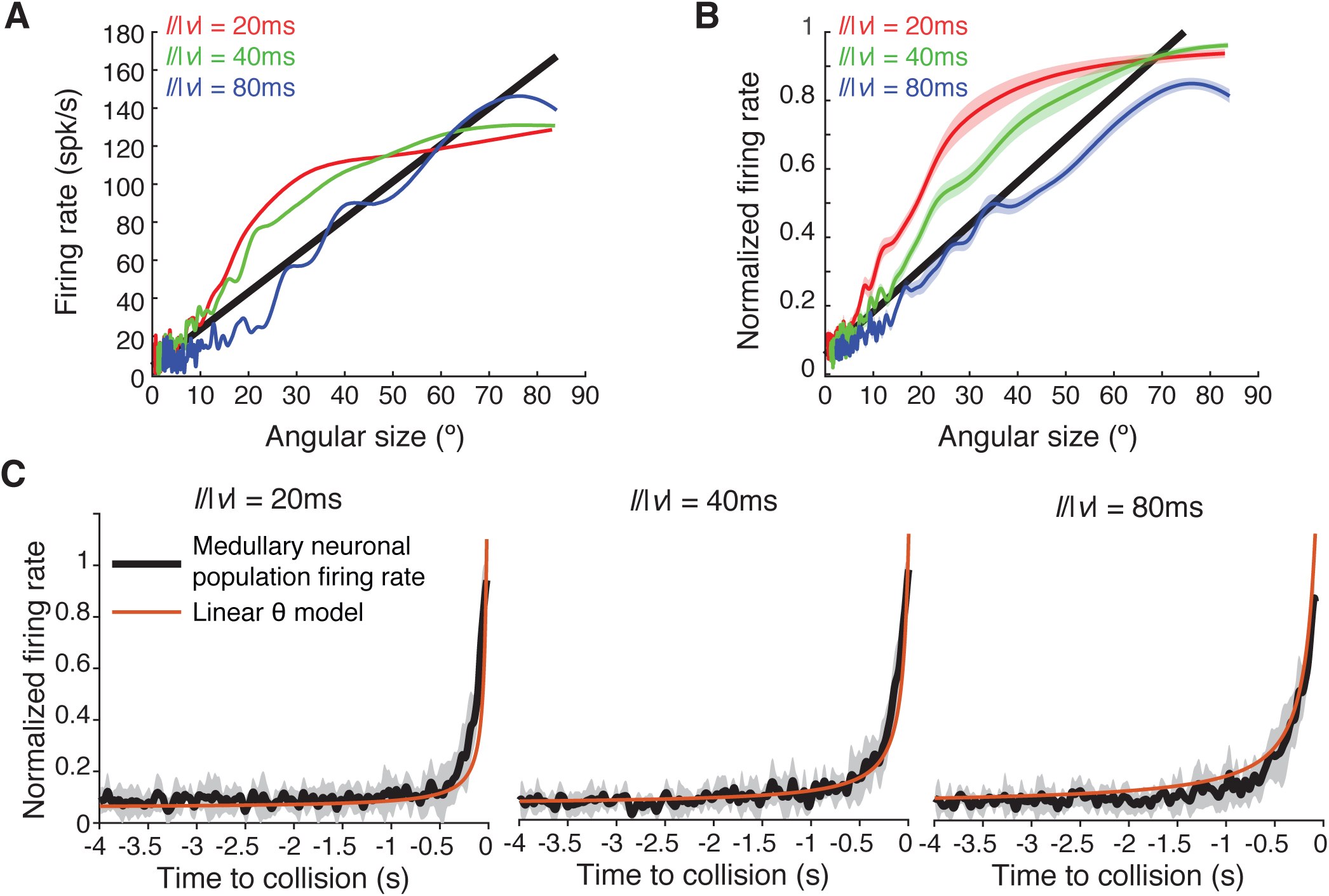
The instantaneous firing rate of medullary DUB neurons presynaptic to the LGMD encodes the angular size of looming stimuli. (A) The IFR of DUB neurons as a function of angular size for looming stimuli at three *l*/|*v*| values (colored lines) and linear fit (black line R^2^ = 0.85, p < 1·10^−9^, F-test) in one example animal (data from other animals in Figure S7). (B) After normalizing firing rates to each animal’s peak firing rate, the linear model was fit to the population average (mean ± s.e.m., n = 6; R^2^ = 0.71, p < 1·10^−9^, F-test). Colors as in (A). (C) Average response during looming approach (black lines and gray shadings are mean ± s.d., n = 6 locusts) with overlaid predicted responses from the linear model in (B) (red lines). For all three *l*/|*v*| values the linear model predicts the time course of the medullary firing (R^2^ = 0.79, 0.93, and 0.90 for *l*/|*v*| = 20, 40 and 80 ms respectively). See also Figure S7.

To test this hypothesis, for each animal we fitted linear models depending on angular size, angular speed or both variables. We found that angular size models well-described the DUB firing (R^2^ = 0.76 ± 0.06) and for all animals performed better than models of angular speed (R^2^ = 0.56 ± 0.05, p = 0.01 rank-sum test). Models using both size and speed provided only a slight improvement over size alone (mean R^2^ = 0.76 vs 0.78, p = 0.62 rank-sum test). We also tested models of squared angular size and speed and log size and speed, but in all cases their performance was again less than that of angular size (squared size: R^2^ = 0.60 ± 0.08, p = 0.01; squared speed: R^2^ = 0.19 ± 0.03, p = 0.0006; log size: R^2^ = 0.42 ± 0.11, p = 0.001; log speed: R^2^ = 0.53 ± 0.10, p = 0.004; rank-sum tests). Additionally, a power law model of angular size did not significantly improve on the linear model (the adjusted R^2^ increased by 0.01) with the best fit power being 1.05 ± 0.27 (mean ± s.d. across animals). As the peak firing rates differed between animals (Figure S7C), we normalized firing rates to each animal’s peak rate and tested the consistency of these findings across animals. The resulting population model (Figure 6B) shows that the angular size dependence of the firing rates is consistent across DUB neurons at all *l*/|*v*| values (R^2^ = 0.71). However, systematic differences from the angular size model in Figure 6B for different l/|v| values suggest that DUB neurons could also code for additional stimulus variables such as angular speed. Finally, the time course of firing was estimated from the model shown in Figure 6B for each *l*/|*v*| value (Figure 6C). In all cases the linear angular size model closely tracked the angular size of the stimulus (R^2^= 0.79, 0.93, and 0.90 for *l*/|*v*| = 20, 40, and 80 ms). These results support the notion that feed-forward inhibition mediated by DUB neurons presynaptic to the LGMD primarily encodes the angular size of the looming stimulus.

### Angular size is encoded by a small, instantaneous firing rate population code

The large receptive fields of individual DUB neurons mean that single cells could accurately encode the angular size of approaching objects. Additionally, the high degree of receptive field overlap and response similarity means that there could be high redundancy in the information conveyed by these neurons. To characterize the population encoding of angular size, we measured both the spontaneous and stimulus evoked spike correlations and tested the angular size encoding accuracy with different numbers of DUB neurons. For this analysis, each pair of simultaneously recorded units was analyzed independently for each trial, yielding 126 sample pairs from 97 stimulus trials (15 medullary neurons, 7 animals).

If the DUB neurons presynaptic to the LGMD had non-overlapping receptive fields, one would expect negative correlations in their looming-evoked responses, as they would be activated at different times during the stimulus. However, since their receptive fields largely overlap, positive spike correlations are predicted. Indeed, during looming stimuli, spike rate correlations averaged 0.28 with 106 of 126 pairs positively correlated (p = 3.2·10^−16^, signed-rank test; Figure 7A, ordinate). To test whether these correlations were due to shared inputs, we also measured the spike rate correlations of the same pairs during spontaneous activity. Surprisingly, this revealed negative correlations for 80 of the 126 pairs (p = 4.6·10^−4^, signed-rank test; Figure 7A, abscissa). This suggests that the stimulus-evoked correlations are due to the overlapping receptive fields, but probably not to the statistics of their shared inputs. Firing rates less dependent on shared input statistics would reduce the redundancy of the information these neurons convey to the LGMD. To test how many DUB neurons are needed to encode angular size during looming, we treated each trial of each unit as independent (yielding sample population sizes of 13-45 pairs per animal from 7 animals; 126 total) and used bootstrap analysis to repeatedly sample different numbers of cells from the population and test their summed firing as a linear predictor of angular size (Figure 7B). For each animal, a small number of units are sufficient to reliably encode angular size (median of 5 cells to reach R^2^ ≥ 0.8). Sampling from more units improved performance to an average peak R^2^ = 0.91±0.04 with less than 10 units needed to reach 95% of peak encoding for 6 out of 7 animals.

**Figure 7.**
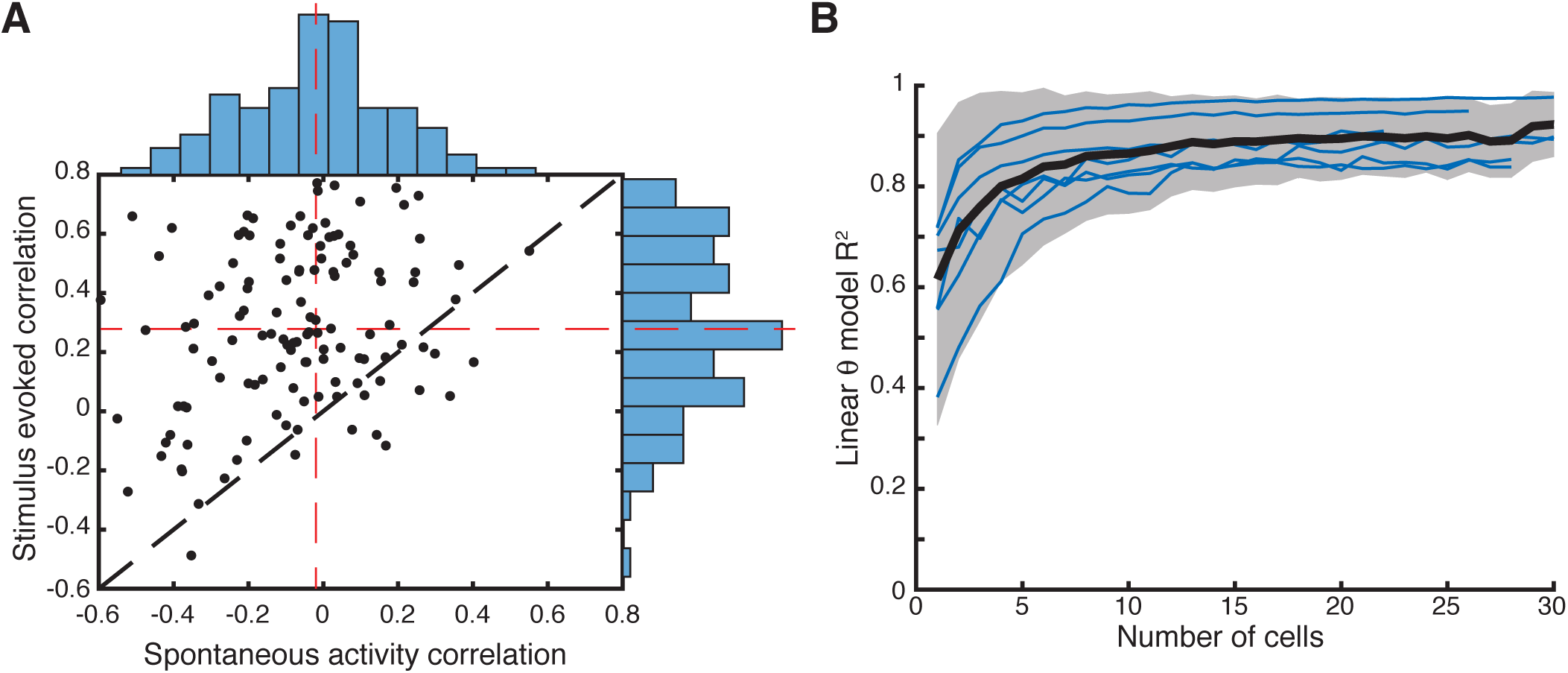
A small population of medullary DUB neurons is sufficient to convey the angular size of looming stimuli. (A) Spike rate correlations between pairs of medullary neurons reveal slightly negative, significant average correlations of spontaneous activity (−0.07 ± 0.22, mean ± s.d., with 80 of 126 pairs negatively correlated, p = 4.6·10^−4^ signed-rank test). During looming stimuli, firing rate correlations increased to 0.28 ± 0.27, mean ± s.d., with 106 of 126 pairs positively correlated (p = 3.2·10^−16^, signed-rank test). Bars are histograms along each axis, with mean correlation values indicated by red dashed lines. (B) Bootstrap estimates of the number of medullary neurons needed to decode angular size. For each animal (blue lines) angular size could be estimated accurately with a small number of neurons, 5.3 ± 3.3 (mean ± s.e.) to reach an R^2^ ≥ 0.8. Black lines and gray shaded region show the mean ± 1 s.d. of the 7 animals.

## Discussion

Feed-forward inhibition is omnipresent in neural circuits and known to play multiple roles related to the temporal processing of sensory information. Here, we focused on a scarcely studied issue: how feed-forward inhibition encodes time-dependent sensory stimulus variables and the role this plays in subsequent downstream dendritic computations. In this study, we characterized the properties of DUB neurons providing feed-forward inhibitory inputs to the LGMD. These neurons have wide receptive fields and a firing rate that depends on the size and speed of moving edges. In response to looming stimuli, their firing pattern resembles that of the LGMD, gradually increasing and peaking before the time of collision. Over its rising phase, the firing rate of DUB neurons is linearly related to the angular size of the looming stimulus, suggesting that they encode as a small, homogeneous population the angular size of approaching objects. Current and earlier results [21] indicate that DUB neurons terminate the excitation triggered in the LGMD by looming stimuli through GABA_A_ receptor gated chloride channels.

### Identifying presynaptic LGMD neurons through *V*_*m*_ spike-triggered averaging

Simultaneous extracellular and intracellular recordings identified DUB neurons whose spikes elicited primarily transient hyperpolarizations and visually evoked IPSPs in the LGMD (Figure 2). The timing of the spike-triggered hyperpolarizations relative to DUB neuron spikes was on the order of 0.05-0.1 ms on average, with a spread of ∼0.3 ms. This delay, the temporal extent and the size of the hyperpolarizations are consistent with those of hyperpolarizing junction potentials caused by gap junctions, as observed in other systems [32]. Although no effective invertebrate gap-junction blocker is presently known and carbenoxolone proved ineffective at blocking the spike-triggered hyperpolarization, a preponderance of evidence points to a mixed electrical and chemical synapse between DUB afferents and the LGMD. Notably, a similar synapse has been described between the LGMD and the DCMD [16, 37, 38].

### Mechanisms of DUB-mediated feed-forward inhibition in the LGMD

Locally puffing GABA near dendritic field C significantly reduced the number of LGMD spikes elicited by looming stimuli. Conversely, puffing picrotoxin onto dendritic field C increased and prolonged responses to the same stimuli [20, 21]. Picrotoxin also abolished spike-triggered visual IPSPs from DUB neurons. In earlier work, immuno-gold tagged GABA antibody staining showed the absence of GABA receptors on dendritic field A, suggesting that GABA-mediated inhibition is localized in dendritic fields B and C [39]. Our results show that electrical stimulation of the DUB induces IPSPs in the LGMD. Conversely, lesioning the DUB abolishes inhibition to OFF stimuli [18a]. Thus, a large body of evidence suggests that feed-forward inhibition to dendritic field C of the LGMD is mediated by GABA_A_ gated chloride channels activated through DUB axons.

### Wide receptive fields of presynaptic inhibitory neurons

Up to now, it was thought that the DUB conveyed inhibitory inputs to dendritic field C through ∼500 axons originating from neurons whose cell bodies lie in the proximal medulla [31]. Based on anatomical considerations, their receptive fields were expected to cover ∼8×12 ° [18a, 30]. Our results, however, do not support this hypothesis. The DUB neurons presynaptic to the LGMD we characterized had wide receptive fields, at least on the order of ∼40×40 °. Furthermore, only a small number of them was required to account for the angular size dependence of feed-forward inhibition. Thus, the DUB neurons presynaptic to the LGMD characterized here build a population much smaller than the ∼500 neurons characterized earlier. We conclude that there must be at least two distinct populations of neurons within the DUB: the small-numbered population we studied electrophysiologically, and the larger population evidenced anatomically.

Although we could record simultaneously only 2 or 3 DUB neurons presynaptic to the LGMD, evidence suggests that their sampling was not random. For instance, the neuron yielding the largest spikes in the extracellular recordings yielded the largest spike-triggered membrane potential hyperpolarization in the LGMD (Figure S2E). This suggests that we were recording from the same neuron across animals. In *Drosophila*, the properties of two types of wide-field excitatory medullary neurons have been recently described: the Lawf2 neurons provide suppressive feedback to large monopolar cells in the lamina [40] and the Tm9 neurons are required for directional motion detection in a broad range of conditions [41]; one recently identified wide-field inhibitory neuron (CT1) has been proposed to modulate the gain of motion detecting T4 cells [42]. Some wide-field inhibitory lobula plate tangential cell (LPTC) characterized in the blowfly, both spiking and non-spiking, such as Vi, VCH and DCH are sensitive to optic flow, but little is known about how their time-varying output influences downstream neurons [43-45].

### DUB neurons presynaptic to the LGMD encode angular size during looming

The firing rate of the LGMD can be described by a multiplication of the angular speed of an approaching object (*θ′(t)*) and a negative exponential of its angular size (exp(*-αθ(t)*)) through a logarithmic-exponential transformation,

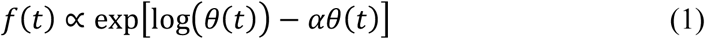

[15, 20, 22]. This formula suggested that the second, inhibitory term could arise from angular-size dependent inhibition provided by DUB neurons. Our experiments confirm this hypothesis. Notably, previous simulations predicted that the population response of DUB neurons would encode the squared angular size, *θ(t)*^*2*^, instead of *θ(t)* [27]. In that work, we had assumed that feed-forward inhibition is mediated by a large population of independent inhibitory neurons with small receptive fields, based on the assumption that they corresponded to the population of DUB neurons characterized anatomically. Under these assumptions, each neuron transmits information about local luminance changes over the area of its receptive field and thus at the population level over the angular area covered by the stimulus. The net result of this spatio-temporal integration process yields an inhibitory input proportional to *θ(t)^2^*. In contrast, the current results show that the size-dependent signals originating from individual ommatidia are not integrated within the small population DUB neurons, but rather lead to an instantaneous firing rate code proportional to *θ(t)*. The encoding of angular size by DUB neurons and the tight correlation between their peak firing rate and that of the LGMD agrees well with results showing that block of feed-forward inhibition terminates the LGMD response to looming stimuli [21]. This has in turn consequences for behavior, because by collectively encoding angular size the DUB neurons ensure that the timing of the LGMD firing rate peak and the subsequent firing rate decay occur when the stimulus reaches a threshold angular size on the retina, as predicted by eq. (1) [22]. Behavioral experiments showed that the firing rate peak is then followed after a fixed delay by the triggering of escape jumps in response to looming stimuli [23, 46].

In other systems, looming-sensitive neurons with response properties resembling those of the LGMD have been described [46-53], but little is known about how inhibitory neurons code for the parameters of an approaching stimulus. In *Drosophila*, the responses of a recently characterized class of looming sensitive neurons (LPLC2) are sharpened by intrinsic inhibitory neurons of the lobula plate (LPi4-3) [54]. Additionally, recent results have evidenced the existence of an angular size-dependent inhibitory input to the giant fiber, a neuron also involved in visually guided collision avoidance behaviors [55]. The neurons mediating this inhibition remain to be identified and may also encode detailed time-varying angular size information. It is thus possible that in this system and in others where feed-forward inhibition plays a role in timing neural responses to sensory stimuli, the neurons mediating feed-forward inhibition also carry detailed time-varying sensory stimulus information. This would confer to feed-forward inhibition a role on par with that of feed-forward excitation in shaping neuronal computations.

## Acknowledgements

We thank Dr. Krapp and Dr. Jiang for comments on the manuscript. The work was supported by grants from the National Science Foundation (DMS-1120952), the National Institutes of Health (NIH) (MH-065339) and NEI Core Grant for Vision Research (EY-002520-37).

## Author Contributions

H.W., R.B.D. and F.G. conceived the experiments; H.W. performed electrophysiology experiments; H.W., R.B.D. and Y.Z. analyzed data; H.W., R.B.D. and F.G. prepared figures and wrote the paper.

## STAR Methods

### CONTACT FOR REAGENT AND RESOURCE SHARING

Further information and requests for resources and reagents should be directed to and will be fulfilled by the Lead Contact, Fabrizio Gabbiani.

### EXPERIMENTAL MODEL AND SUBJECT DETAILS

Wild type locusts (*Schistocerca americana*) were maintained under a 12:12 h light:dark life cycle at a mean temperature of 31°C and fed with wheat seedlings and oat meal. All experiments were performed on eight to ten week-old adult locusts of either sexes.

### METHOD DETAILS

#### Animal preparation

The dissection was carried out as described previously [20]. Briefly, the legs, wings and antennas were removed. The locust body was mounted in a plastic holder with the dorsal side facing upward. The head capsule was opened and the muscles above the optic lobes and brain were removed. The air sacs above the ventral nerve cords were removed to expose them. During the dissection, the whole head was immersed in ice-cold locust saline solution. The right eye was fixed with bee wax to the holder. The entire procedure lasted approximately 1.5 h. During experiments, the head remained immersed in saline, except for the right eye which was exposed to the stimulus-displaying monitor.

#### Electrophysiology

Thin-walled borosilicate glass capillaries (1.2/0.9 mm outer/inner diameter; WPI, Sarasota, FL) were used for making intracellular electrodes to record from the LGMD. A Sutter P-97 Flaming/Brown type micropipette puller (Novato, CA) was used to fabricate sharp electrodes which were filled with an internal solution containing 1.5 M KAc, 1M KCl and 10 mM Alexa 594 (final micropipette resistance: 8 – 20 MΩ). Bath solutions included 140 mM NaCl, 5 mM KCl, 5 mM MgCl_2_, 5 mM CaCl_2_, 6.3 mM HEPES, and 4 mM NaHCO_3_, and 73 mM Sucrose. The pH was adjusted to 7.0. All the intracellular recordings were recorded in bridge mode using a NPI amplifier (NPI electronics, Tamm, Germany). The electrode resistance and capacitance were compensated before approaching the surface of the optic lobe. The LGMD and the descending contralateral movement detector (DCMD) have a 1:1 spiking correspondence [12]. The LGMD was thus identified using DCMD spikes recorded extracellularly from the ventral nerve cord with a pair of stainless steel hook electrodes. The spikes of the DCMD were the biggest ones recorded extracellularly and could therefore easily be identified. The LGMD was initially stained with Alexa 594 by iontophoresis (- 2 nA current pulses, alternating between on and off every second). Next, the sharp electrode was inserted into the thin dendrites of dendritic field C receiving feed-forward inhibitory inputs under visual control. Initially, sharp tungsten electrodes with an input impedance of 0.5 and 5 MΩ (FHC, Bowdoin, ME) were compared to record spikes from the medulla. Use of the lower impedance tungsten electrode resulted in a better yield, with a larger number of isolated units. Therefore, in the following experiments we mainly used a pair of sharp tungsten electrodes with an input impedance of 0.5 MΩ. In five experiments, we used electrodes with an input impedance of 2 MΩ, which yielded similar results. The electrodes were positioned in the DUB at the level of the proximal medulla, near the inhibitory branches of the LGMD to record the extracellular spikes generated by medullary DUB neurons. The distance between the paired tungsten electrodes amounted to 50-75 μm and their distance to the intracellular electrode positioned in dendritic field C of the LGMD was at least 200 µm. Earlier paired recordings in the optic lobe have shown that extracellular spikes can be recorded up to 100 µm away from a neuron impaled using an intracellular electrode, with no noticeable interference between the recordings [56]. The location of the DUB was identified by electrical stimulation with an amplitude and duration varied from 15 to 320 µA and from 0.1 to 1 ms through the sharp tungsten electrodes using a stimulus isolator (DS3, Digitimer Ltd., Fort Lauderdale, FL) which produced characteristic IPSPs in the LGMD (Figure 1D). The extracellularly recorded DCMD and medullary spikes were amplified through a differential AC amplifier (A-M Systems 1700; Sequim, WA) and an instrument amplifier (Brownlee 440; NeuroPhase, Santa Clara, CA). All the recordings were sampled at 19927.5 Hz. A chlorided silver wire was placed in the bath solution and used as a reference electrode.

#### Visual stimulation

Looming stimuli, moving edge, OFF flash, luminance change and grating stimuli were generated using custom software with a computer running the QNX4 real-time operating system (QNX, Ottawa, ON) and displayed through a cathode ray tube monitor with a 200 Hz refresh rate. The right eye was positioned to face the center of the monitor from a distance of 20 cm. In these experiments, looming stimuli refer to black disks on a white screen expanding rapidly in size from a tiny point at the center of the screen, simulating the approach of an object with a half-size *l* and a constant translation speed *v* [21, 22]. The time course of approach is characterized by the ratio *l*/|*v*|. Three *l*/|*v*| ratios were used, *l*/|*v*| = 20, 40 and 80 ms. Looming stimuli were presented every other 4 minutes. Moving edge stimuli were used to map the spatial receptive field of the medullary DUB neurons. In these experiments, a black edge on a white screen moved either from posterior to anterior or from ventral to dorsal, and vice versa, with a constant speed. The width of the edge was one–fourth (or one-eighth) of the screen dimension perpendicular to the edge movement direction. Thus, when the edge moved dorso-ventrally its width was 18.8°; when the edge moved antero-posteriorly, its width was 22.8°. The edge speed took the following six values: 7.1, 14.2, 28.4, 56.8, 113.6, and 227.2 °/s to measure its relationship with the medullary DUB neurons’ firing rate. A speed of 28.4 °/s was used to map the spatial receptive fields, with additional experiments at 14.3°/s. Edge stimuli were presented at an interval of 5 s and were cycled through all positions and motion directions in turn to minimize potential habituation of the medullary neurons’ firing rate. Local grating stimuli consisted of 3 alternating white and black bars. They had a width and height equal to 1/3 of the screen size, and moved in one direction at a speed of 28.4 °/s. During a trial, each grating stimulus first appeared on the screen and remained stationary for 10 s, then started moving over the next 10 s, and finally disappeared. OFF Flash stimuli consisted of 4 black squares with size set to 8.6, 17.2, 34.4 and 68.8°. Luminance changes from white to black varied linearly from 68.5 to 0.4 cd/m^2^ over 6 durations of 0.1, 0.2, 0.5,

1, 2, and 5 s.

#### Injection of picrotoxin and GABA in the lobula

To study the pharmacological properties of feed-forward inhibition, picrotoxin (5 mM) and GABA (10 mM) dissolved in water were puffed along the dorsal edge in the lobula, close to the region where the inhibitory dendrites of LGMD’s field C arborize. The injected solution contained 0.5 % of the colorant fast green to visualize the tip of the injection pipette and the amount of solution injected in the lobula. The injection pipette tip diameter varied between 1 and 2.5 µm. After injection, the dye diffused around the injection site and stayed confined to the lobula. A picospritzer was used to control the duration and the puffing pressure (8 psi/55 kPa; WPI, Sarasota, FL). Based on earlier work [18], the estimated final concentration of drugs at the level of field C was ≤ 200 µM (picrotoxin) and ≤ 300 µM (GABA).

### QUANTIFICATION AND STATISTICAL ANALYSIS

Custom Matlab code was used for data analysis. The medullary DUB neuronal spikes were sorted following general principles exposed in [58], using a spike-sorting program modified from publicly available code described in [58]. Briefly, the data were first normalized, before detecting spikes with a threshold and building an event dictionary consisting of non-overlapping spike waveforms. After performing principal component analysis to reduce dimensionality, the data set was visualized using the grand tour visualization technique implemented in the software GGobi. Typically, individual spike clouds were well separated and could be clustered using K-means classification based on the visualization step. Next, the delay of individual spike events relative to the spike template for a given cluster was estimated and cancelled, before subtracting from the original recording each event identified as being from that cluster. This last step of the procedure [59] was repeated until no further spikes could be identified in the original recording. In particular, this procedure allows to efficiently resolve most cases of spike superposition. Because the extracellular and intracellular recordings were acquired at a sampling rate of 20 kHz, our measurements of the timing of the peak *V*_*m*_ hyperpolarization relative to the presynaptic DUB spikes is limited to a resolution of ± 0.1 ms in Figures 2D and E. In a few cases (8.6%), we observed slightly smaller delays of approximately −0.2 ms. These recordings were excluded from the subsequent data analysis. To calculate instantaneous firing rates (IFRs) during looming stimuli, the spike trains of the LGMD and the medullary DUB neurons were convolved with a Gaussian filter having a standard deviation of 20 ms. The medullary neurons’ IFR evoked by edge or grating movement, luminance decrements, and OFF flashes was calculated as briefly explained below (see [60] for details). For each spike train, the firing rate before the time of the first spike was set to 0 (*f(t)* = *0* if *t*<*t*_*1*_, where *1* is the index of the first spike); the firing rate at the time of the first spike was calculated as half the inverse of the time interval between the first and second spike (*f*(*t*) = 0.5*/*(*t*_*2*_*-t*_*1*_) if *t* = *t*_*1*_); then, the firing rate at a time between two neighboring spikes was calculated as the inverse of the corresponding inter-spike interval (ISI, *f*(*t*) = 1*/*(*t*_*i*_*-t*_*i-1*_) if *t*_*i-1*_<*t*<*t*_*i*_); the firing rate at the time of subsequent spikes was calculated as the average of the inverse inter-spike intervals immediately preceding and following it (*f*(*t*) = 0.5*/*(*t*_*i*_*-t*_*i-1*_) + 0.5*/*(*t*_*i+1*_*-t*_*i*_) if *t* = *t*_*i*_); the firing rate at the time of the last spike was half of the inverse of the last ISI (*f*(*t*) = 0.5/(*t*_*n*_*-t*_*n-1*_) if *t* = *t*_*n*_, where *n* is the index of the last spike); the firing rate after the time of the last spike was set to 0 (*f*(*t*) = 0 if *t*>*t*_*n*_). Above, *t*_*1*_ and *t*_*n*_ refer to the first and last spike time, *f*(*t*) is the IFR at a certain time *t*, and *i* indexes the spikes. To map visual responses to specific stimulus locations on the screen in Figure 3, we estimated the neural response delay by considering two responses to a pair of visual stimuli moving in opposite directions. We inverted in time one of these responses and computed the temporal shift needed to align it with the other one. For this purpose, the two mean IFRs averaged across trials were first smoothed over a one second window using the Matlab function ‘filtfilt’ which implement zero-phase digital filtering. The peak firing rate was computed for each of the two traces and then the two intervals spanning the range were the firing rate was over ¾ of the peak. The temporal shift was estimated as half of the offset required to align the center of these windows. In figure 3C, we computed the mean and standard deviation of the spatial firing rate distribution along the azimuth and elevation axes by treating it as a two-dimensional probability distribution after normalization.

The membrane hyperpolarization evoked by electrical stimulation was calculated as the difference between the LGMD mean *V*_*m*_ within 0.1 s before the artifact caused by the electrical stimulus and the minimum *V*_*m*_ within 0.02 s after the electrical stimulus artifact. Because electrical stimulation always elicited initially excitation in the LGMD we could only estimate an upper bound in the onset latency of inhibition. This latency of inhibition, which we call the earliest detectable latency, was calculated as the difference from the time of the electrical stimulation artifact to the subsequent time when LGMD *V*_*m*_ returned to the mean *V*_*m*_ value computed within 0.1 s before the artifact, as explained above.

The Wilcoxon signed-rank test was used to compare the statistical differences between groups treated with or without picrotoxin or GABA. Unless denoted with mean ± s.e. (standard error), all the data were described as mean ± s.d. (standard deviation).

For linear modeling of the data presented in Figure 6 and the corresponding text, we used the Matlab function ‘fitlm’ which generates a best fit from a least-squares linear regression. Simple linear models with constant and linear terms (no interaction terms) were employed using the summed firing rates of the medullary neurons. The peak firing rate of individual medullary neurons and the number of neurons recorded both varied across animals. So, before we averaged data from multiple animals (Figures 6D, E), the firing rates for each animal were normalized to the peak firing rate of that animal. The quality of the fit for each model was measured by the adjusted-R^2^ statistic and the F-statistic of the model’s linear component. For comparisons between models (e.g., *θ*(*t*) and *θ′*(*t*)) a Wilcoxon rank-sum (Mann-Whitney U) test was used based for each animal on the ajusted-R^2^ model values.

Analyses of spike correlations (Figure 7) were conducted among pairs of units recorded in the same stimulus trial. If 5 stimulus trials were presented to an animal in which 3 medullary DUB units were recorded, this yielded 15 sample pairs (5 comparisons between units 1 and 2, 5 between units 1 and 3, and 5 between units 2 and 3). In total 126 paired comparisons were made from 15 medullary units recorded from 7 animals. For the spontaneous correlations, the firing of each sample pair just before the start of the visual stimulus was analyzed (2.1 ± 1.0 s of recording with average spontaneous firing rates of 24 ± 13 spk/s). The data were down-sampled to 200 Hz and instantaneous firing rates were calculated with Gaussian convolution as described above.

To determine how many medullary DUB neurons were needed to accurately predict the stimulus angular size, we binned the data and used a bootstrap sampling method. The time periods corresponding to 1° increments of the stimulus angular size were determined for each stimulus trial (in total 209 trials from 7 animals) and the average IFR of individual medullary units was measured within each corresponding time period. This was done to facilitate combining responses to stimuli with different *l/|v|* values. Once the data was binned according to the stimulus angle that evoked it for each trial, 1000 bootstrap samples (using Matlab ‘bootstrap’ function) were taken with different numbers of cells ranging from 1 to the population size for that animal. For each of the 1000 draws for each sample size, the firing rates of the drawn units were summed together and fit to the stimulus size. The average quality of those fits (R^2^) is displayed in Figure 7B.

### DATA AND SOFTWARE AVAILABILITY

Data and Matlab code to recreate all figures available at doi:10.17632/h5y2pphkvp.1

